# SFRP1 shapes astrocyte to microglia cross-talk in acute and chronic neuroinflammation

**DOI:** 10.1101/2020.03.10.982579

**Authors:** Javier Rueda-Carrasco, María Inés Mateo, Aldo Borroto, María Jesús Martin-Bermejo, Markus P. Kummer, Stephanie Schwartz, José P. López-Atalaya, Balbino Alarcon, Michael T. Heneka, Pilar Esteve, Paola Bovolenta

**Affiliations:** Centro de Biología Molecular Severo Ochoa, CSIC-UAM, c/ Nicolás Cabrera, 1, Campus de la Universidad Autónoma de Madrid, Madrid 28049, Spain; CIBER de Enfermedades Raras (CIBERER) Madrid 28049, Spain; Universitätsklinikum Bonn, Neurology; German Center for Neurodegenerative Diseases (DZNE) 53127 Bonn, Germany; Instituto de Neurociencias, CSIC-UMH, Sant Joan d’Alacant, Spain

**Keywords:** Neuroinflammation, microglia, HIF pathway, LPS, EAE, reactive astrocytes, Multiple Sclerosis, Alzheimer’s disease

## Abstract

Neuroinflammation is a common feature of many neurodegenerative diseases, which fosters a dysfunctional neuron-microglia-astrocyte crosstalk that, in turn, maintains microglial cells into a perniciously reactive state that often enhance neuronal damage. The molecular components that mediate this critical communication are however not fully explored. Here, we have asked whether Secreted-Frizzled-Related-Protein-1 (SFRP1), a multifunctional regulator of cell-to-cell communication, is part of the cellular crosstalk underlying neuroinflammation. We show that in mouse models of acute and chronic neuroinflammation, astrocyte-derived SFRP1 is sufficient to promote and sustain microglial activation, and thus a chronic inflammatory state. SFRP1 allows the upregulation of components of Hypoxia Induced Factors-dependent inflammatory pathway and, to a lower extent, of those downstream of the Nuclear Factor-kappaB. We thus propose that SFRP1 acts as a critical astrocyte to microglia amplifier of neuroinflammation, representing a potential valuable therapeutic target for counteracting the harmful effect of chronic inflammation present in several neurodegenerative diseases.

## Introduction

Degeneration of neurons occurs in a variety of rare and common pathological conditions of the CNS including, for example, Alzheimer’s disease (AD) or Multiple Sclerosis (MS). CNS function is supported by a robust neurons-glia crosstalk [1,2], so that neuronal damage is almost invariably associated with the activation of two types of glial cells: microglia and astrocytes. The prompt glial response to insults brings about an inflammatory reaction that favors tissue healing and helps restoring CNS homeostasis [3,4]. However, excessive glial activation, as often occurs in neurodegeneration, is itself a cause of neuronal loss, which, in turn, establishes a state of pernicious chronic neuroinflammation [4–7]. Consistent with an important cellular crosstalk, glial cell dysfunction, either as a consequence of hyper-or hypo-functionality, can also be the cause of neuronal damage, rather than its consequence, strongly contributing to the progression of neurodegenerative diseases [3,4,8,9]. Although this is nowadays a widely accepted idea, there is still only partial information on the molecular components that sustain a dys-or hyper-functional state of glial cells and an abnormal glia/neuron crosstalk. Here we have investigated whether SFRP1 may represent one of such components.

SFRP1 is a small, secreted and dispersible protein with a dual function: modulation of Wnt signaling [10] and inhibition of ADAM10 activity [11]. The latter function is particularly relevant in the context of neurodegeneration because ADAM10, a member of the A Disintegrin and Metalloprotease family of plasma-membrane proteins [12], acts as an α– secretase (or sheddase; [13] for many substrates expressed in both neurons and glial cells. In microglial cells, ADAM10-mediated shedding of TREM2 (Triggering Receptor Expressed on Myeloid) is a relevant mechanism for controlling their activation [14]. In neurons, ADAM10 activity releases the ectodomain of CD200 and CX_3_CL1 [15,16], two components of the mechanism by which neurons maintain microglial cells in a surveying state [2]. Consistent with these roles and the additional regulation of proteins involved in synapse formation [12,17], genetic inactivation of *Adam10* in mice causes neuroinflammation and loss of synaptic plasticity [18]. Interest in ADAM10 is further increased by genetic studies supporting a link between impaired ADAM10 activity and AD [19,20]. Thus, and based on its non-amyloidogenic processing of the Amyloid Precursor Protein (APP) [21], ADAM10 may have a protective role against AD [22]. Consistent with this possibility, we have recently shown that elevated levels of SFRP1, its endogenous negative modulator, contribute to AD pathogenesis [23]. SFRP1 upregulation correlates with poor ADAM10-mediated processing of APP and of the synaptic protein N-Cadherin, whereas neutralization of its activity prevents the appearance of AD pathological traits, including glial cell activation [23].

*Sfrp1* is abundantly expressed in mammalian radial glial progenitors of the developing CNS [11,24,25] but is largely down-regulated in the adult brain with the exception of restricted neurogenic areas [25–27]. Besides in human neurodegenerative diseases [23,28–30], in which the mRNA localizes to glial cells [23,27], SFRP1 upregulated expression has been reported in several other inflammatory conditions such as periodontitis, rheumatoid arthritis, uropathies or pulmonary emphysema [10,31]. An upregulation has been observed also in the aged brain [29], in which a low-grade chronic inflammation is present [32]. Notwithstanding, whether SFRP1 is directly involved in the modulation of neuroinflammation remains unexplored.

By addressing this issue, here we show that Sfrp1 is a novel mediator of the astrocyte to microglia crosstalk that underlies mammalian CNS inflammation. In mice, astrocyte-derived SFRP1 is sufficient to activate microglial cells and to amplify their response to distinct acute and chronic neuroinflammatory challenges, sustaining their chronic activation. From a molecular point of view, SFRP1 allows for the full expression of down-stream targets of the transcription factors hypoxia induced factors (HIF) and, to a lesser extent, nuclear factor-kappa B (NF-κB), which are mediators of neuroinflammatory responses [33,34]. Thus, neutralizing SFRP1 function may represent a strategy to counteract pernicious chronic neuroinflammation that contributes to many human neurodegenerative conditions.

## Results

### Acute brain neuroinflammation elevates astrocyte-specific levels of SFRP1 expression

To evaluate a possible link between SFRP1 and neuroinflammation, we induced acute brain inflammation by injecting bacterial lipopolysaccharides (LPS; [35] or control saline into the somatosensory cortex of 3 months-old *Sfrp1*^*+/βgal*^;*CX*_*3*_*CR1*^*+/GFP*^ mice (n=4), which allow the simultaneous identification of microglial (GFP; [36] and *Sfrp1* producing cells (nuclear βgal-positive; [37]. Immunofluorescence of the brains three days after injection, when inflammation is at its peak [38], revealed a broader βgal immunoreactivity (reporter of *Sfrp1* expression) in LPS vs saline treated animals (Fig. 1A), largely localized in GFAP^+^ astrocytes but not in Iba1^+^ microglial cells (Fig. 1A; S1A). Immunodetection of SFRP1 with specific antibodies [23] confirmed an increased SFRP1 production after LPS compared to saline injections (Fig. 1B). Notably, SFRP1 protein distribution in saline injected animals was comparable to that of non-injected mouse brains [23]. Consistent with the secreted and dispersible nature of SFRP1 [39,40], immuno-signal was widely distributed in the brain parenchyma (arrows, Fig. 1B). The use of a highly specific ELISA [23] further confirmed a significant increase of SFRP1 levels in extracts of small cortical tissue samples from LPS treated animals (Fig. 1C) as compared to that present in equivalent brain regions from non-injected or saline injected animals (Fig 1C). Together these data show that astrocytes produce and secrete increased level of SFRP1 in response to a bacterial lipopolysaccharide.

**Figure 1.**
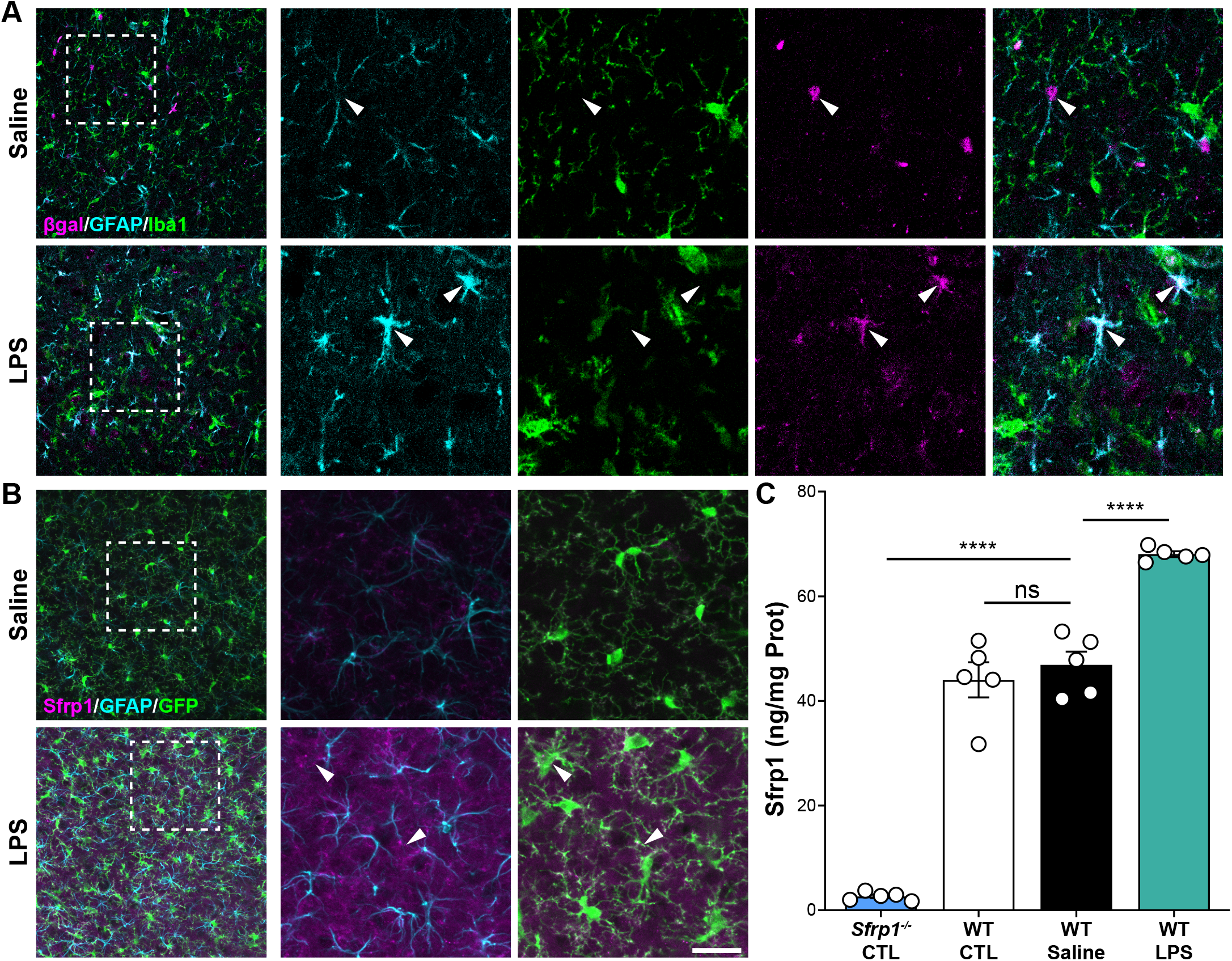
Astrocytes upregulate SFRP1 expression upon LPS stimulation. A, B) Confocal image analysis of cryostat sections from adult *CX*_*3*_*CR1*^*+/GFP*^;*Sfrp1*^*+/βgal*^ mouse brains three days after intra-cortical infusion of saline or LPS. Sections were immunostained for βgal (magenta) and Iba1 (green, A) or Sfrp1 (magenta) and GFP (green, B), and GFAP (cyan). Arrowheads indicate βgal/GFAP (A) and Sfrp1/GFP co-localization (B). Scale bar 25μm. C) ELISA determination of Sfrp1 levels in brain extracts from 3 months-old wt and *Sfrp1*^*-/*-^ mice. WT mice were injected either with saline or LPS. Three days after injection the region around the injected side (10 mm^3^ cortical cube) was isolated and SFRP1 content compared with that present in similar region of non-injected or *Sfrp1* null mice used as negative control (n=5 mice for each group). Error bars represent Standard Error. Statistical significance: ****P<0.0001; One-way ANOVA followed by Bonferroni test.

### In vivo Sfrp1 gene addition is sufficient to trigger and sustain glial cell activation

We next reasoned that if SFRP1 is indeed associated with neuroinflammation, its forced expression should be sufficient to activate glial cells. To test this possibility, we infused lentiviral particles (LV) containing *Sfrp1*-IRES-*Gfp* or control IRES-*Gfp* into the lateral ventricle of 10 weeks-old wt mice (n=8; Fig. 2). As expected by the injection site, GFP^+^ (used to determine infection efficiency) LV-transduced cells were largely found lining along the wall of the lateral ventricle, in the choroid plexus and, to a less extent, along the rostral migratory stream (Fig. S1B). Immunohistochemistry of cortical sections 1 month after injection showed a significantly higher presence of SFRP1 protein at the infected site (Fig 2A) associated with GFAP^+^ reactive astrocytes and Iba1^+^ microglial cells as compared to LV-IRES-*Gfp* control animals (Fig. 2A,B; quantifications performed in 3 of the injected animals). Iba1 immunoreactivity was accumulated around the injection site (Fig. 2A,B) and many cells had a round amoeboid morphology and a significantly higher CD45 expression (Fig. 2B; S1C), two characteristics of reactive microglia [41]. CD45^hi^ round cells could also represent blood-borne infiltrating macrophages (Fig. S1C), although the molecular and functional difference between the two cell types is currently a matter of debate [42]. A wider distribution of hyper-phosphorylated Tau, which often appears as a brain response to inflammation and degeneration [43], was also detected in LV-*Sfrp1*-IRES-*Gfp* transduced vs control brains with a significantly wider distribution (Fig. 2A,B). Analysis of a different set of LV-transduced animals (n=4) 5 months after LV delivery showed a persistent microglial activation in SFRP1-vs GFP-treated animals, with more CD45^+^ cells than those observed at 1-month post-injection (Fig. S1C). Furthermore, several CD45^hi^-round-shaped microglia/macrophages were detected in the parenchyma, especially in the proximity of transduced cells (white arrowheads in Fig. S1C). Together these results indicate that SFRP1 is sufficient to trigger an inflammatory response of glial cells, which persists for prolonged periods of time.

**Figure 2.**
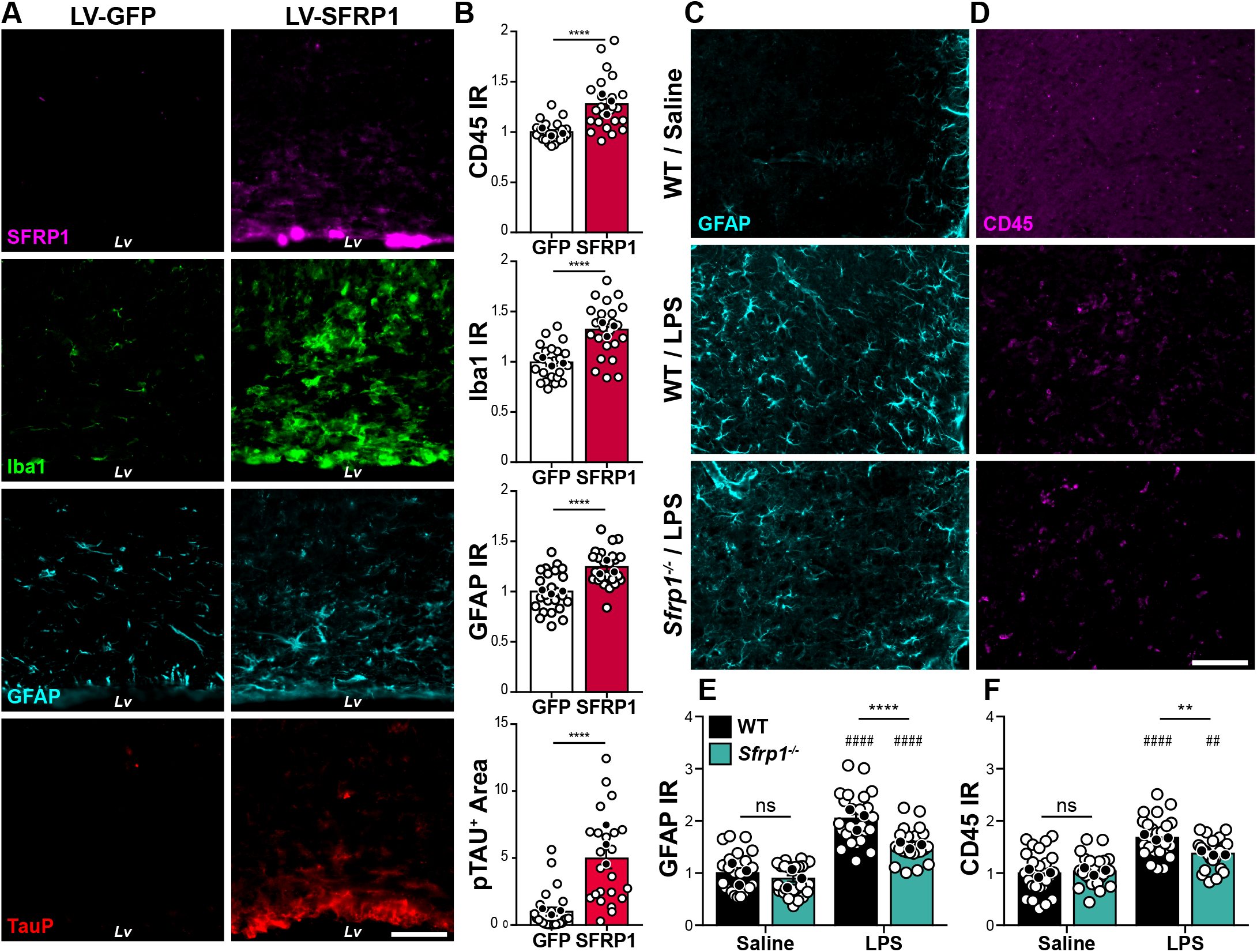
SFRP1 is sufficient and required to enhance glial cell activation upon LPS treatment. A) Coronal cryostat sections of LV-IRES-*Gfp* or LV-*Sfrp1*-IRES-*Gfp* infected brains 1-month post-infusion, immunostained for SFRP1 (magenta) Iba1 (green) GFAP (cyan), or TauP (red). Scale bar 100μm. B) The graph shows the level of GFAP, Iba1 and CD45 immunoreactivity (IR) and the area occupied by TauP signal (n=24 acquisitions, white dots; N=3 mice, black dots, for each group), normalized to LV*-IRES-Gfp* infected brains. Error bars represent Standard Error. Statistical significance: ****P<0.0001 by two-sided Student’s t-test. C, D) Coronal sections from wt and *Sfrp1*^*-/-*^ mouse brains three days after infusion of saline or LPS, immunostained for GFAP (cyan, A) or CD45 (magenta, B). The images are high power views of the somatosensory cortex (lower power view in Fig S2). Scale bar 60μm. E, F) The graphs show the levels of immunoreactivity (IR) for GFAP (C) and CD45 (D, P=0,006) present in cortical sections (n=24 acquisitions white dots; from N=3 animals, black dots, per group) from wt and *Sfrp1*^*-/-*^ animals infused with saline or LPS. Bars represent Standard Error. Statistical significance: ** or ^##^ P<0.01, **** or ^####^ P<0.0001 by two-way ANOVA followed by Bonferroni’s multiple comparisons test. * and # indicate significance between genotypes and treatments, respectively.

### Sfrp1 is required for amplifying CNS inflammatory response

Given that SFRP1 was sufficient to induce glial cell activation, we next asked whether it was also a necessary component of the CNS inflammatory response. To this end, we took advantage of *Sfrp1*^-/-^ mice, in which SFRP1 protein is completely undetectable [23] (Fig 1C). These mice have a slightly shorter and thicker cortex that however does not affect their life span, reproduction rate or cognitive and motor behavior and present no evident neuronal defects [23,25]. Furthermore, their content of astrocytes and microglial cells was undistinguishable from that of wt mice (Fig. 2C-F; Fig S1D). We thus compared the effect of intra-cortical LPS infusion into the brains of 3-months-old wt and *Sfrp1*^-/-^ mice (Fig. 2C,D). In the somatosensory cortex of wt brains (Fig. S1D), LPS but not saline treatment caused the appearance of GFAP^+^ reactive astrocytes (Fig. 2C) and CD45^+^ reactive microglia (Fig. 2D). In contrast, *Sfrp1*^-/-^ littermates presented fewer and less immuno-positive astrocytes and a significant reduction of CD45^+^ reactive microglia/macrophages (Fig. 2C,D), suggesting that SFRP1 is relevant for the activation of both astrocytes and microglial cells. Quantitation of immunoreactivity among different genotypes and treatments confirmed no significant differences between the two saline-treated genotypes but showed a significantly lower response of *Sfrp1*^-/-^ mice to LPS as compared to wt (Fig. 2E,F). Notably however, *Sfrp1*^-/-^ mice do not completely lose their respond to LPS, given that this is significantly different from that observed in saline injected mice (Fig. 2E,F). This suggests that SFRP1 is involved in boosting inflammation.

To evaluate a possible specificity of this response, we induced Experimental Autoimmune Encephalomyelitis (EAE) in wt and *Sfrp1*^-/-^ mice. This is a widely used experimental model for MS, a human inflammatory demyelinating disease. In EAE, CNS inflammation and gliosis occur as a consequence of a strong autoimmune response against the peripheral exposure to myelin components [44], thus representing a neuroinflammatory paradigm quite different from the direct LPS intra-cerebral infusion. Female littermates of the two genotypes were immunized following a standard protocol and animals were scored for the development/remission of their clinical symptoms over the course of a month [45]. Animals were classified with a standard 0 to 5 rank based on their paralysis degree, with 0 corresponding to absence of symptoms and 5 to a moribund condition [45]. In wt mice (n=19), tail limping -the first symptom of the disease-became apparent around 8 days after immunization with a subsequent rapid progression, so that, by 16 days, most of the wt animals presented hind limb paralysis followed by a slow recovery (Fig. 3A). Notably, 47% of the immunized wt mice developed extreme and protracted symptoms (Fig 3A,B). The response of *Sfrp1*^-/-^ mice (n=19) to immunization was instead slower and milder: only 16% of them developed extreme symptoms and their recovery was significantly faster (Fig. 3A,B). Immunostaining of spinal cord sections before immunization showed no differences between wt and *Sfrp1*^-/-^ in astrocytes, microglia or myelin distribution (not shown). In contrast, in animals (n=4) sacrificed 16 days after immunization, there was a significant reduction of pathological signs in *Sfrp1*^-/-^ vs wt mice, including infiltration of CD4^+^ lymphocytes, presence of Iba1^+^ macrophages/activated microglial cells and GFAP+ reactive astrocytes, pial surface disruption and loss of MBP^+^ myelin (Fig. 3C,D).

**Figure 3.**
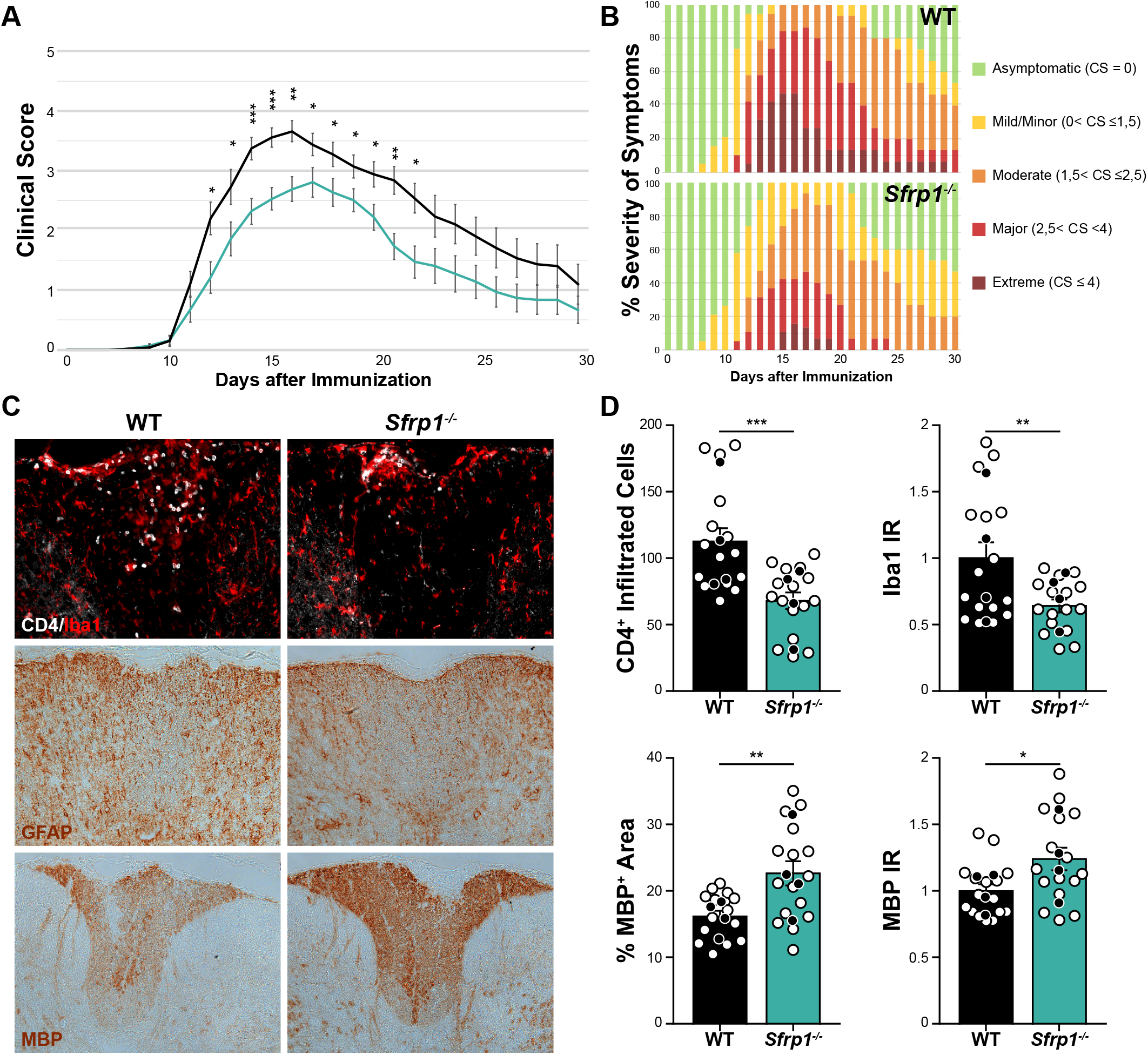
*Sfrp1*^*-/-*^ mice develop a milder form of EAE. A, B) Time course analysis and severity of the symptoms in wt and *Sfrp1*^*-/-*^ mice after EAE induction. In A means are depicted with black (wt) and cyan (*Sfrp1*^*-/-*^) lines. In B data are expressed as % of the total number of analyzed animals (n=19 per genotype). B) Wt and *Sfrp1*^*-/-*^ mice immunized with MOG and sacrificed 16 days post immunization. Cryostat sections of the thoracic spinal cords were stained with antibodies against CD4, Iba1, GFAP and MBP. Images show the region dorsal fasciculus. Scale bar 200μm. C) Quantification of CD4+ infiltrated lymphocytes, Iba1 immunoreactivity and MBP immunoreactive area in the dorsal fasciculus of spinal cord sections from wt and *Sfrp1*^*-/-*^ mice 16 days after immunization (n=16 acquisitions, white dots; from N=4 animals, black dots, per genotype). Statistical significance: *P<0.05, **P<0.01, ***P<0.001 by Kolmogorov-Smirnov test followed by Mann–Whitney U nonparametric test comparing mice of same day after immunization (a) or two-sided Student’s t-test (c).

All in all, these data indicate that SFRP1 is commonly required for a robust neuroinflammatory response, likely mediating astrocyte to microglial crosstalk.

### Astrocyte-derived Sfrp1 is required for a robust response of microglial cells to damage

The latter possibility found support in SFRP1 localization around microglial cells (Fig. 1B) and in the remarkably poor microglial activation (Fig. 2; Fig. 3) and myeloid cell recruitment (Fig. 3C) observed in the absence of *Sfrp1*. To further explore this possibility we used flow cytometry analysis to determine the proportion of the different CD11b^+^ myeloid populations [46] in the cortex of *CX*_*3*_*CR1*^*+/GFP*^ and *CX*_*3*_*CR1*^*+/GFP*^;*Sfrp1*^*-/-*^ mice 3 days after intra-cerebro-ventricular LPS or saline administration (Fig. 4A; Fig S2A). In the absence of *Sfrp1*, there was a significant decrease of LPS-induced infiltration of CD11b^+^/CD45^+^/GFP^-^ monocytes and of the proportion of microglial cells that passed from a CD11b^+^/CD45^lo^/GFP^+^ surveying to a CD11b^+^/CD45^+^/ GFP^+^ activated state (Fig. 4A,B). Furthermore, FACS analysis of isolated GFP-positive microglial cells (Fig. S3A) revealed that, upon infusion of pHrodo *E*.*coli* bioparticles [47] into the third ventricle, microglial cells derived from *CX*_*3*_*CR1*^*+/GFP*^;*Sfrp1*^*-/-*^ mouse brains showed less cumulative phagocytized fluorescent signal than those isolated from *CX*_*3*_*CR1*^*+/GFP*^ brains at both 40 and 65 hr post infusion (Fig. 4C; Fig. S3). Notably, at 40 hr the percentage of phagocyting microglial cells in *CX*_*3*_*CR1*^*+/GFP*^;*Sfrp1*^*-/-*^ mice was significantly higher than in controls (Fig. 4C) but this proportion underwent a drastic reduction at 65 hr (Fig. 4C). These data suggest that in SFRP1 absence microglial cells present a less efficient phagocyting activity that, at earlier time points, might be compensated by increasing cell recruitment. A differential rate of phagocytosis vs degradation or the acidity of the lysosomal compartment [48] may explain the time course of phagocytosis observed in mutant microglia.

**Figure 4.**
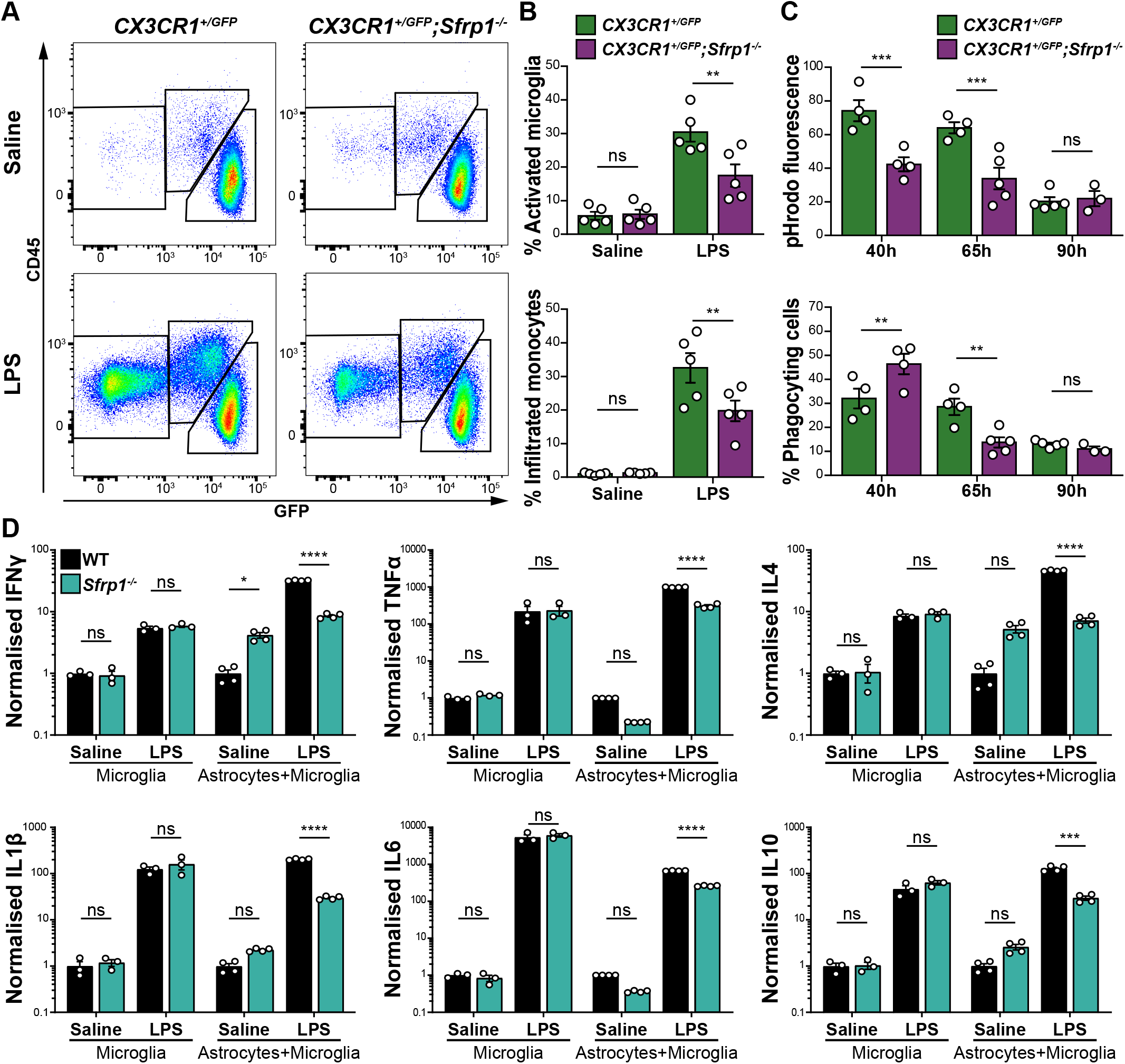
SFRP1 amplifies LPS-induced microglial activation and enhances their cytokine secretion. A) Representative cytometry plots of CD11b+ populations present in the cortex of *CX*_*3*_*CR1*^*GFP/+*^ or *CX*_*3*_*CR1*^*GFP/+*^;*Sfrp1*^*-/-*^ mice 3 days after saline or LPS intraventricular infusion. Gating was set for isolating the following myeloid cell populations: CD11b^+^;CD45^lo^;GFP^+^ surveying microglia, CD11b^+^;CD45^+^;GFP^+^ activated microglia and CD11b^+^;CD45^+^;GFP^-^ infiltrated monocytes. B) Quantification of the percentage of activated microglia and infiltrated monocyte present in *Sfrp1*^*-/-*^ and controls brains 3 days after infusion of saline or LPS. C) Quantification of the total fluorescence from phagocytized pHrodo-labelled *E. coli* bioparticles and percentage of phagocyting cells in the CD11b^+^;GFP^+^ microglial population. Error bars represent Standard Error. Statistical significance: **P<0.01, ***P<0.001 by two-way ANOVA followed by Bonferroni’ s multiple comparisons test. D) Secretory profile of cytokines present in the medium of primary microglia or microglia and astrocytes cultures from wt and *Sfrp1*^*-/-*^ pups exposed for 24h to saline or LPS (1μg/ml). The content of IFNγ, TNFα, IL1β, IL4, IL6 and IL10 present in the culture media was determined by ELISA. Values, normalized to those of untreated wt cultures, are represented in Log scale. Error bars represent Standard Error. Statistical significance: ns P>0.05; *P<0.05; ***P<0.001; ****P<0.0001 by two-way ANOVA followed by Bonferroni’s multiple comparisons test.

Collectively, these data support the contention that SFRP1, mainly derived from astrocytes, modulates microglial activation in response to an inflammatory challenge and favors their efficient function. To assess that indeed there is an astrocyte-to-microglial flux of information, we next established microglial or astrocyte/microglia mixed cultures (1:1 ratio) from wt and *Sfrp1*^*-/-*^ neonatal mice and used a multiplex ELISA to determine their cytokines’ release (IFNγ, TNFα, IL1-β, IL4, IL6 and IL10) in the culture medium, as a measure of their LPS-induced activation (Heneka et al., 2014). At 24 hr, LPS but not saline treatment enhanced the accumulation of all tested cytokines in the culture media of purified microglial cultures with no significant differences between genotypes (Fig. 4D). In astrocytes/microglia cultures derived from *Sfrp1*^*-/-*^ LPS-induced cytokines’ release was instead significantly reduced, as compared to that detected in similar wt-derived cultures (Fig. 4D). ELISA determination confirmed the presence of SFRP1 only in culture media of astrocytes/microglia mixed cultures derived from wt pups (Fig. S2B), strongly supporting that astrocyte-derived SFRP1 enhances microglial activation.

### Sfrp1 allows for the full expression of down-stream targets of HIF transcription factors

We next reasoned that if SFRP1 enhances microglial activation, its absence should modify their transcriptomic profiling towards a less activated or surveying state. To test this possibility, we used fluorescence-associated cell sorting to purify GFP^+^ microglial cells from the brain of *CX3CR1*^*+/GFP*^ and *CX3CR1*^*+/GFP*^;*Sfrp1*^*-/-*^ mice (n=4). Cells were purified three days after the injection of LPS or saline and used to obtain the corresponding transcriptomic profiles. We initially compared the gene-expression signature of *CX3CR1*^*+/GFP*^ and *CX3CR1*^*+/GFP*^;*Sfrp1*^*-/-*^derived microglial cells in response to LPS (Fig. 5). Principal-component analysis demonstrated LPS as the main source of variation (72% variance), whereas only 6% of the total variations could be attributed to the genotype after secondary-component analysis (Fig. 5A, Table S1A-D), well in line with the observation that SFRP1 is poorly expressed in the adult brain under homeostatic conditions (Fig. 1; Esteve et al., 2019b). The relatively central position of LPS-treated *Sfrp1*^*-/-*^ microglia supported their milder inflammatory response (Fig. 2, 3), which was further confirmed when overall gene expression variations of LPS-treated control and *Sfrp1*^-/-^ microglia were plotted and compared (Fig. 5B,C). Linear regression analysis of this comparison demonstrated a 30% reduction of the global transcriptomic response induced by LPS in the absence of *Sfrp1* (slope of red dotted line, Fig. 5D). LPS treatment in control microglia induced the upregulation of 1128 genes, 487 of which were shared with *Sfrp1*^-/-^ microglia (Table S1A-B). LPS treatment also induced the upregulation of 121 and the down-regulation of 167 microglial genes in the absence of SFRP1 (Fig. 5E; Table S1A-D). Genes associated with metabolic pathways (i.e. *AldoA, LdhA* or *Pygl)*, the cell cycle (i.e. *Gas6*) or immune regulators (i.e. *Mif, Mefv*) were highly upregulated in controls but not in *Sfrp1*^*-/-*^ microglia (Table S1A-D), whereas genes downstream of the Toll-like 4 receptor (TLR4), such as *Tlr2*, were upregulated at similar levels in both genotypes (Table S1A-D). TLR4 is fundamental for LPS recognition and the activation of the immediate inflammatory response (Ransohoff and Perry, 2009), strongly suggesting that, in the absence of *Sfrp1*, microglial cells retain their prompt response to damage, as also shown by their cytokines’ release upon LPS treatment (Fig. 4D).

**Figure 5.**
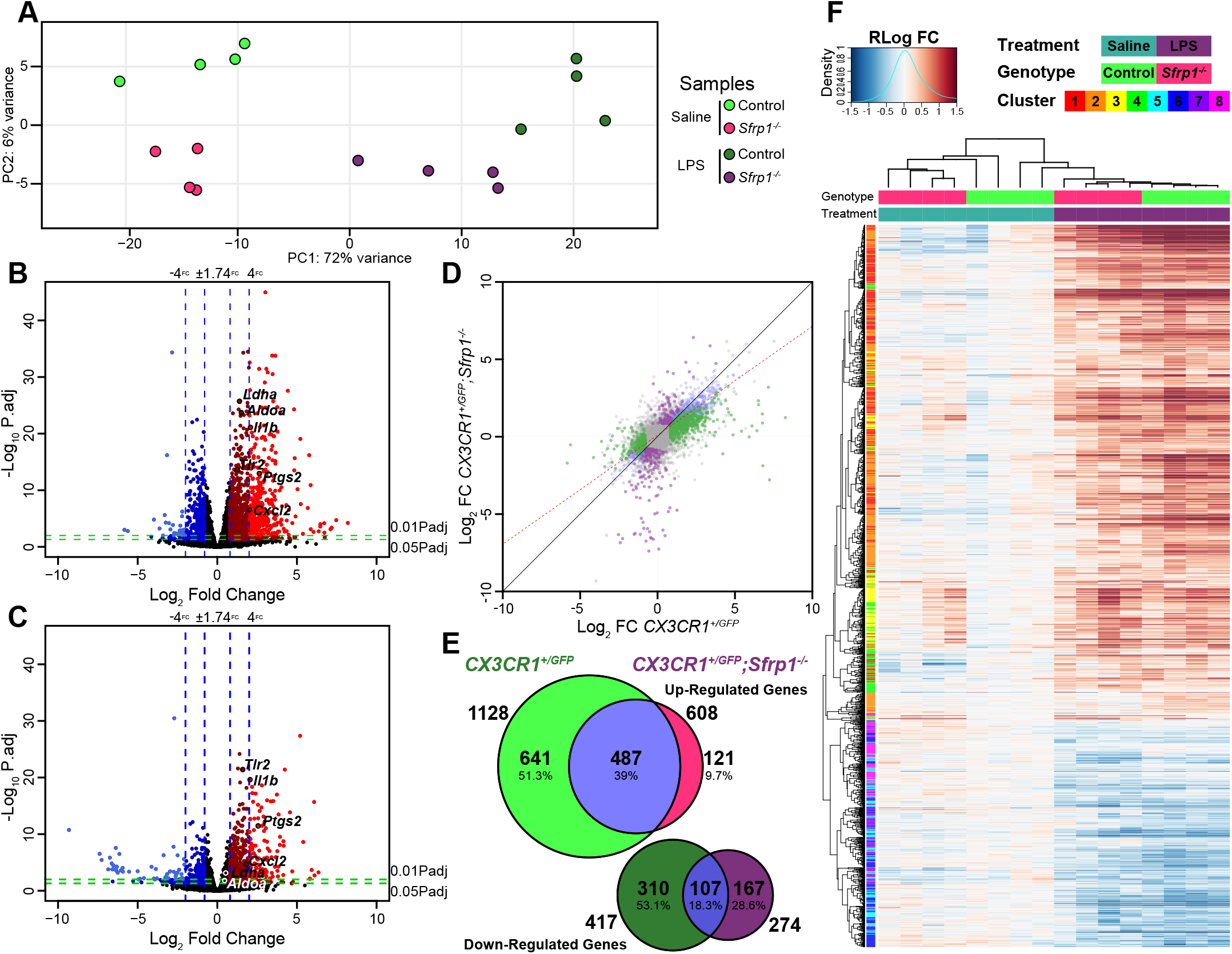
SFRP1 enhances the transcriptional response of microglial cells to LPS treatment. A) Principal component analysis with the 1000 most variable genes from microglial cells isolated from *CX*_*3*_*CR1*^*GFP/+*^ and *CX*_*3*_*CR1*^*GFP/+*^;*Sfrp1*^*-/-*^ mouse brains three days after saline or LPS intra-cerebro-ventricular infusion. The analysis depicts sample clusterization by genotype and treatment. B, C) Volcano plots of differential gene expression from *CX*_*3*_*CR1*^*GFP/+*^ (B) or *CX*_*3*_*CR1*^*GFP/+*^;*Sfrp1*^*-/-*^ (C) microglial cell in response to LPS. Data are represented as Log2 Fold Change vs −Log10 adjusted p-value. Blue vertical lines represent an increase of 75 and 400% in the expression levels respectively. Green horizontal lines represent a 0.05 or 0.01 adjusted p-value of statistical significance respectively. D) Scatterplot of differential expressed genes in *CX*_*3*_*CR1*^*GFP/+*^ and *CX*_*3*_*CR1*^*GFP/+*^;*Sfrp1*^*-/-*^ microglia; the red line represents linear regression (slope 0.704014 p-value: < 2.2e-16) of the differential expression, indicating response attenuation. Colored dots represent response which are exclusive for each genotype as represented in E. E) Venn diagram showing the extent of differential gene expression between *CX*_*3*_*CR1*^*GFP/+*^ and *CX*_*3*_*CR1*^*GFP/+*^;*Sfrp1*^*-/-*^ microglial cells. Fold change (75%) and adjusted p-value < 0.05 cut-off. F) Heatmap showing fold changes of regularized log transformed gene-level RNA-seq counts with hierarchical clustering of samples and differentially expressed genes

Consistent with this observation, hierarchical clustering of the samples demonstrated treatment-dependent similarities, but with *Sfrp1*^-/-^ microglia showing an attenuated response to LPS treatment as compared to wt (Fig. 5F). Hierarchical clustering of the differentially expressed genes presented a pattern of covariance that was further analysed by Z-score covariance unsupervised clustering (Fig. 5F). This analysis generated 4 different clusters of upregulated genes, which were analysed for enrichment of regulatory elements and gene ontology annotations (Fig. S4). Genes regulated by E2F transcription factors and implicated in cell cycle regulation composed the less abundant 3 and 4 clusters. Genes belonging to cluster 3, involved in the regulation of chromosomal segregation and spindle organization, were slightly more upregulated in control microglia. On the contrary, genes belonging to cluster 4 and involved in chromatin assembly and DNA packing were upregulated in *Sfrp1*^-/-^ microglia, suggesting possible differences in the length of the cell cycle (Fig. S4).

Cluster 1 and 2 were the largest clusters and included genes that mediate the inflammatory response and regulators of the defense response (Fig. S4A,B). Both clusters were strongly upregulated in control microglia in response to LPS (Fig. S4A): genes in cluster 1 showed significant enrichment in NF-kB TF binding sites at the promoter regions, whereas those putatively controlled by Hif transcription factors were enriched in cluster 2 (Fig. S4C; Table S1E,F). In microglia derived from *Sfrp1*^*-/-*^ mice, the expression of a few genes belonging to cluster 1 was decreased (Fig. S5A; Table S6), whereas that of genes belonging to cluster 2 was basically abrogated (Fig. S4A; Table S1F). Ingenuity Pathway Analysis network representation of the NF-κB and HIF downstream targets showed that the absence of SFRP1 has a greater effect on downstream targets of HIF (Fig. 6A). Genes downstream NF-κB, such as *Tlr2* and *ApoE* showed LPS-dependent up-regulation with levels almost similar in the two genotypes, whereas the expression of the homeostatic *Cx3cr1* and *Trem2* genes was unchanged (Fig. 6B; Table S1E). Of note, a number of genes shared between the NF-κB and HIF pathways, such as *Cxcl2, Cxcl3* and *Ptgs2*, were strongly up-regulated in LPS-treated control microglia but not in those derived from *Sfrp1*^*-/-*^ brains (Fig. 6A). Similarly, other LPS-induced HIF targets failed to be up-regulated in the absence of *Sfrp1* and maintained levels of expression comparable to those of saline-treated control microglia (Fig. 6A,B; Table S1F). These included *Vegfa, Sod2, Mif, Aldoa* and *Ldha*. Notably, *Hif1α* expression was similar to that observed in LPS treated controls (Table S1F), possibly in line with the notion that Hif1 levels and activity are largely modulated by post-translational modifications and proteolysis [49].

**Figure 6.**
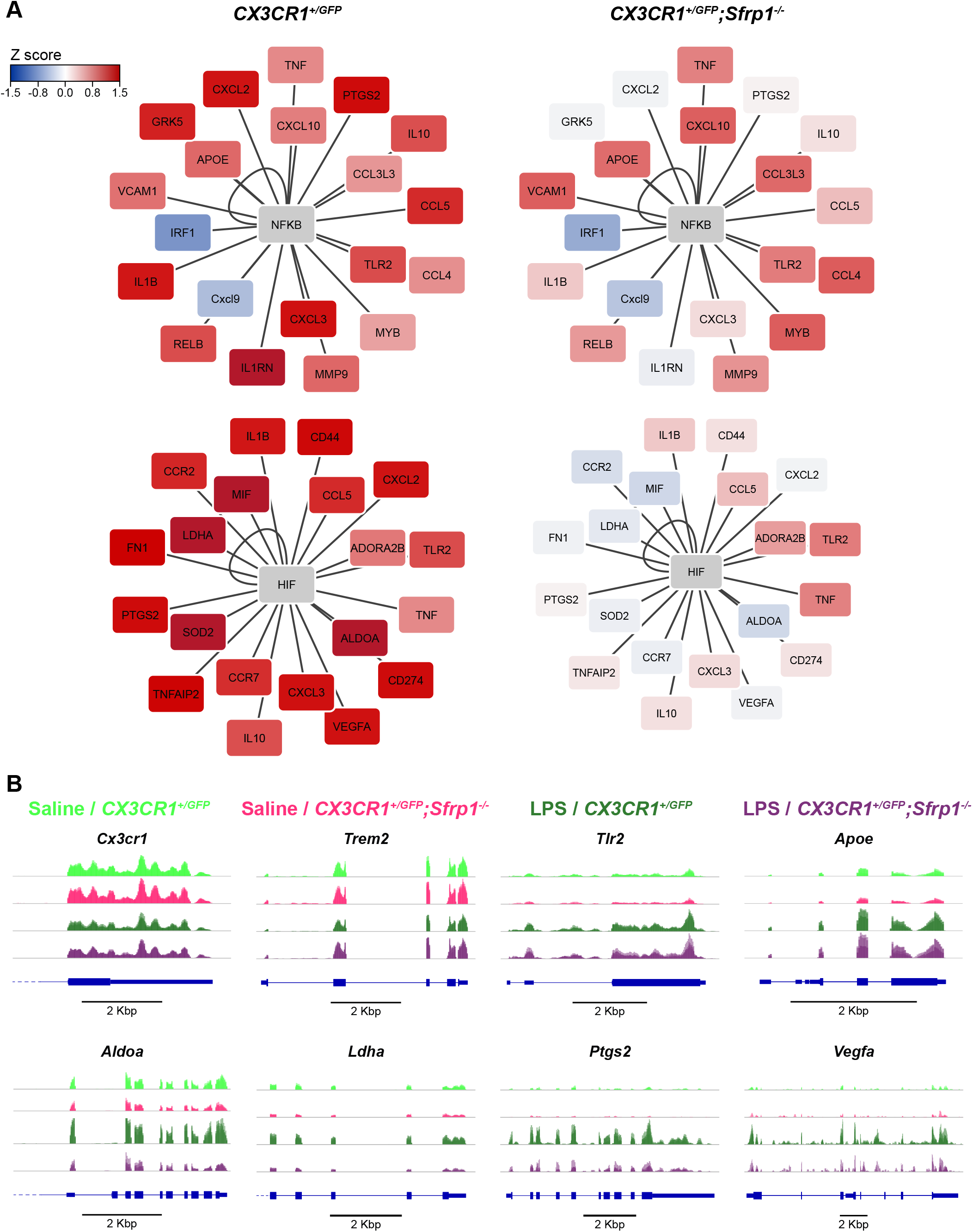
SFRP1 induces the up-regulation of HIF target genes during neuroinflammation. A) Ingenuity Pathway Analysis network representation of NF-κB and HIF downstream targets for *CX*_*3*_*CR1*^*GFP/+*^ and *CX*_*3*_*CR1*^*GFP/+*^;*Sfrp1*^*-/-*^ microglial cells. Colored by Z-score alteration of their expression after LPS stimulus. B) Integrative Genomics Viewer transcription profile of represented genes for saline and LPS treatments of *CX*_*3*_*CR1*^*GFP/+*^ and *CX*_*3*_*CR1*^*GFP/+*^;*Sfrp1*^*-/-*^ microglial cells. Scale bar: 2 Kbp.

All in all, these results support the idea that in the absence of *Sfrp1* microglial cells can sense and respond to an inflammatory insult undertaking an initial response. SFRP1 is however required to enhance this response allowing for the full activation of microglial-mediated inflammation in the brain.

## Discussion

Microglial cells transit from a surveying to an activated state in response to adverse signals derived from the surrounding environment [50]. However, both insufficient or prolonged microglia activation is harmful to the brain so that an elaborated neuron-microglia-astrocyte cross talk is in charge of shaping the brain immune response to damage [51]. The molecular components that mediate the flux of information from neurons to microglia (i.e. the CD200/ CD200R signaling system) [50] or from microglia to astrocytes (i.e. cytokines and NO) [1] have been in part identified. In contrast, there is perhaps less information on how astrocytes communicate with microglial cells [1]. Our study unveils that SFRP1 is part of the molecular signals that astrocytes provide to microglial cells to enhance their inflammatory response. In response to damage, reactive astrocytes produce and secreted SFRP1, which, in turn, increases the number of activated microglial cells and fosters the expression of HIF and, to a much lower extent, NF-kB down-stream targets in microglial cells. SFRP1 persistency is however pernicious as it sustains a chronic inflammatory state. Indeed, its inactivation significantly reduces the prolonged neuroinflammation associated with EAE. These observations indicate that SFRP1 is a potential valuable target to counteract the harmful effect of prolonged inflammatory conditions, such as those present in MS or AD.

SFRP1 function has been studied in many developmental contexts largely linked to the control of cell specification, proliferation and differentiation [52] but its homeostatic role in adult tissues is less clear. Its poor or absent expression (or that of other SFRP family members) due to promoter hypermethylation is considered a sign of bad prognosis in different types of tumor [10,53,54], although *Sfrp1*^*-/-*^ mice are viable, fertile and do not form tumor spontaneously [55], indicating that SFRP1 absence per se is not tumorigenic. On the other hand, our finding unveils that elevation of SFRP1 levels has also pathological consequences but related to neuroinflammation (this study) and neurodegeneration [23]. As an advantage, SFRP1 neutralization in AD-like mice counteracts disease pathology without any apparent side effect [23]. In the specific case of neuroinflammation, neutralization of SFRP1 function should have less effect on the initial acute inflammatory phase, which is a necessary step towards pathogen elimination, tissue repair and homeostasis restoration [51]. Abrogation of SFRP1 function does not prevent this acute response given that, in its absence, both LPS-infusion and MOG-immunization induce an initial inflammatory reaction, albeit at somewhat lower levels. Furthermore, activated microglial cells and infiltrated monocytes can still be isolated from the brain of *Sfrp1*^-/-^ mice, although at significantly decreased numbers. These microglial cells retain phagocytic competence, although with a significant less efficiency and no significant difference was observed in the LPS-induced cytokines release from purified microglial cells obtained from *Sfrp1*^-/-^ and wt brains. This agrees with the observation that in homeostatic conditions SFRP1 is very poorly expressed in the adult mammalian brain [23] and this study). Our transcriptomic analysis shows that *Sfrp1*^-/-^ microglia cells retain their capacity of sensing LPS, providing strong support to the implication of SFRP1 in the amplification of the inflammatory response. Indeed, *Tlr2* expression, a read-out of LPS-mediated TLR4 activation [38], is increased (and to a comparable extent) in microglial cells from both genotypes. A similar consideration applies to a large fraction of the genes related to the NFκB-dependent pathway, which is largely related to the acute inflammatory response [34].

Brain response to LPS, but also to other harmful signals, depends on the fast reaction of functional microglial cells [56], whereas astrocytes and neurons do not initially and directly participate in this response despite their reported expression of the TLR4 receptor [57,58]. Reactive microglial cells thereafter induce astrocytes activation that, in turn, feeds back on microglial function [1]. Fitting our data into this loop, we propose that up-regulation and release of SFRP1 from astrocytes occurs as part of their microglia-mediated early activation. Secreted SFRP1 then acts on microglial cells amplifying a HIF-dependent inflammatory response and thus a significant increase in cytokine secretion, further impinging upon inflammation. Supporting SFRP1 implication in this astrocyte-microglia loop, astrocytes *Sfrp1*^*-/-*^ mice show decreased GFAP expression and thus are also less reactive, likely producing lower levels of other microglia-activating molecules. This implies that, as long as SFRP1 up-regulation persists, neuroinflammation persists, contributing to its chronicization. This idea is well in agreement with our observation that genetic inactivation of *Sfrp1* significantly limits the severity and progression of EAE, a condition that, as MS, is characterized by persistent microglial activation. Both EAE and MS are driven by the infiltration of peripheral macrophages and lymphocytes [59]. We cannot exclude that the lack of peripherally expressed *Sfrp1* may limit this infiltration and thus the disease, especially because SFRP1 has been shown to influence lymphocytes’ differentiation [60]. Notwithstanding there is strong evidence that endogenous microglial activation is critical for EAE onset and maintenance [61,62] and the *SFRP1* gene is hypomethylated in brain samples from MS patients [63], likely promoting an abnormal protein increase. Thus, reduced microglial activation is a plausible cause of the milder EAE symptoms observed in *Sfrp1*^-/-^ mice.

SFRP1 up-regulation in other neurodegenerative diseases characterized by the presence of chronic inflammation further supports a role of SFRP1 microglial activation. An example is Glaucoma [64] or AD [23,30], in which antibody-mediated neutralization of SFRP1 strongly decreases different AD pathological features, including neuroinflammation [23]. Notably, SFRP1-mediated inhibition of ADAM10, a metalloprotease responsible for non-amyloidogenic processing of APP, contributes to the generation of toxic amyloid peptides in AD patients, whereas SFRP1 deficiency decreases amyloid burden [23]. Low neuroinflammation upon SFRP1 neutralization could therefore be secondary to this reduction [23]. The present study shows that this is not necessarily the case as SFRP1 acts directly on microglial cells, suggesting that SFRP1 simultaneously impinges upon multiple pathological events in AD. A possible SFRP1-mediated activation of microglial cells in AD finds further support in our transcriptomic analysis that links SFRP1 with HIF signaling. Indeed, many LPS-induced genes regulated or co-regulated by Hif were not activated in the absence of *Sfrp1*, although *Hif1α* expression was not modified, possibly because its levels are largely post-translationally regulated [49]. Consistent with our observations, a number of recent studies have shown that microglial cells isolated from AD-like mouse models undergo important metabolic changes with the activation of the HIF pathway [65–67]. Furthermore, *Hif1α* is involved in astrocytes proinflammatory activity [68] and its activity seems to be dysregulated in association with other genes genetically linked to AD risk, suggesting that HIF-1α may be detrimental in AD pathology [66].

The precise molecular interactions underlying SFRP1-mediated HIF pathway activation are at the moment an open question. Microglial cells express members of the ADAM family of metalloproteases [14], including ADAM10 which has been shown to be upregulated in both neurons and microglial cells during microglial mediated synaptic pruning [69]. Furthermore, SFRP1 effectively inhibits ADAM enzymatic activity in cultured microglial cells (our own observations). Given that TREM2 is a proven substrate of ADAM10/17 [14], it seems plausible to postulate that SFRP1 may modulate the shedding of TREM2 on the surface of microglial cells. This shedding occurs, for example, in response to LPS induced activation of TLR4 [70,71], thereby attenuating microglial activation [70]. Notably, microglial cells deficient in TREM2 undergo only a partial and abortive activation and remain locked in an almost homeostatic state [72,73]. It is thus tempting to speculate that in SFRP1 absence enhanced shedding of TREM2 may reduce microglial activation as we have observed in *Sfrp1*^*-/-*^ mice, whereas high SFRP1 levels, by interfering with ADAM function, may enhance TREM2 signaling. In addition (or alternatively), SFRP1 may interfere with ADAM10-mediated shedding of the neuronal ligands CX3CL1 and CD200 preventing the generation of their soluble forms [15,16] and thus the activation of their respective microglial receptors, CX3CR1 and CD200R [36,74]. Unfortunately, lack of appropriate biochemical tools has prevented us from verifying these possibilities in our mouse models, leaving this question open for future studies.

An alternative and non-mutually exclusive possibility involves SFRP1 function as a Wnt signaling modulator. The role of this pathway in microglial activation is however somewhat controversial, perhaps in line with the notion that Wnt pathway activity is highly context dependent. For example, studies *in vitro* have shown that Wnt3a and Wnt5a can counteract LPS-induced microglia activation [75]. In contrast, in preterm-born infants and in different postnatal animal models of injured-induced neuroinflammation, downregulation of the expression of Wnt signaling components seems a prerequisite for pro-inflammatory activation of microglia [76]. SFRP1 upregulation could lead to the same net effect, inducing at least the down-regulation of the expression of *Axin2* or *Lef1*, genes that targets and read-out of Wnt/βcatenin pathway activation. Notably however, we found no evidence of changes in the expression of these genes in our RNaseq analysis, making us favor the possibility that SFRP1 acts through ADAM10.

In conclusion, we have shown that astrocyte-derived SFRP1 plays an important role in shaping microglia response to CNS damage, sustaining chronic neuroinflammation. This effect might not be limited to the brain, as SFRP1 upregulation has been reported in different pathological conditions associated with inflammation or fibrosis such as periodontitis, rheumatoid arthritis, uropathies or pulmonary emphysema [10,31]. Therefore, once analyzed the possible side effect, SFRP1 neutralization [23] may represent a promising therapeutic avenue to treat a wide variety of chronic pathological conditions.

## Material and Methods

### Animals

Newborn and adult mice of both sexes were used, unless otherwise indicated. Mice were maintained in pathogen–free conditions at the CBMSO animal facilities, following current national and European guidelines (Directive 2010/63/EU). Experimental procedures were approved by the CBMSO and Comunidad Autónoma de Madrid ethical committees. *Sfrp1*^*-/-*^ mice were generated by inter-cross of the *Sfrp1*^*-/-*^;*Sfrp2*^*+/-*^ mice in a mixed 129/C57BL/6 background as described [11] and then back-crossed at least 4X with C57BL/6J to unify the background. Wild type (wt) animals were littermates selected from heterozygous crosses. Breeding pairs of *CX*_*3*_*CR1::GFP* mice [36] were kindly provided by Prof. J Avila, CBMSO. Mice were further crossed with the *Sfrp1*^*-/-*^ to obtain *CX*_*3*_*CR1::GFP;Sfrp1*^*-/-*^.

### Stereotactic LPS infusion

LPS or saline were infused into the brain parenchyma of 10-12 weeks-old littermates from wt and *Sfrp1*^*-/-*^ or *CX*_*3*_*CR1*^*+/GFP*^ and *CX*_*3*_*CR1*^*+/GFP*^;*Sfrp1*^*-/-*^ mice. Animals were anaesthetized with 4% Isoflurane (Forane, AbbVie Farmacéutica) vaporized into a sealed anesthetic induction chamber (SurgiVet, Smiths Medical) and placed in a stereotaxic apparatus (Stoelting). Anesthesia was maintained at 2.5% in 250ml/min oxygen flow. Saline (2.5µl) or LPS (5µg; Escherichia coli 0111:B4; Sigma Aldrich) were delivered through a small skull window using a Quintessential Stereotaxic Injector (Stoelting) coupled to a 10µl syringe with a 34G needle (Hamilton) at the rate of 0.5µl/min using the following bregma coordinates: 0.0mm A-P; −1.0 mm lateral, and −1.5mm D/V. Mice were let survive for three days and then sacrificed and processed for further analysis. Delivery of pHrodo Red *E*.*coli* BioParticles Conjugate (Molecular Probes) was performed with a similar procedure at - 2.5 mm A-P; 0.0 mm lateral, and −2.3 mm D-V from bregma. Lentiviral vectors (2.5μl), generated as described below, were delivered 0.5 mm A-P; 1.0 mm lateral and −2.3 mm D-V from bregma. Mice were let survive 1-5 months and then analyzed.

### Lentiviral (LV) particle generation

LV particles carrying IRES-*Gfp* or *Sfrp1-*IRES-*Gfp* were obtained by transient transfection of mycoplasma-free HEK-293T cells [77,78]. Cells were transfected employing a three-plasmids HIV-derived and VSV pseudotyped LV system kindly provided by M.K. Collins, University College London, UK; A. Thrasher, Institute of Child Health, UK; and D. Trono, Ecole Polytechnique Fédérale de Lausanne, Switzerland. Culture supernatants were collected two and three days after transfection and ultra-centrifuged. Pellets containing the concentrated particles were resuspended in PBS. Functional titter of the viral preparations was determined by transfection of HEK-293T cells (1 × 10^8^ TU/ml, in both cases).

### Experimental Autoimmune Encephalomyelitis (EAE)

Chronic EAE was induced as described [45]. Briefly, 8-10 weeks-old female C57BL/6J and *Sfrp1*^*-/-*^ littermates were injected bilaterally in subcutaneous femoral region with 150 mg of MOG35–55 (Espikem) emulsified with Freund’s complete adjuvant (Sigma Aldrich), supplemented with Mycobacterium tuberculosis (1mg/ml; H37Ra strain from Difco), followed by two intraperitoneal injections of pertussis toxin (200 ng; Sigma Aldrich) separated by 48h. An observer blind to the animals’ genotype weighed and inspected the animals daily to detect the appearance of clinical signs according to the following classification: 0) no overt signs of disease; 1) weakness at the tail distal portion; 1.5) complete tail flaccidity; 2) moderate hind limb weakness; 2.5) severe hind limb weakness; 3) ataxia; 3.5) partial hind limb paralysis; 4) complete hind limb paralysis; 4.5) complete hind limb paralysis with muscle stiffness; 5) moribund state and hence sacrificed according to ethical procedures. A representative pool of mice was anesthetized and perfused intracardially sixteen days after immunization, when symptoms picked, for histological analysis. Other animals were let recover. Spinal cords were dissected and processed.

### Primary cultures

Glial primary cultures were established from cerebral cortices of C57BL/6J or *Sfrp1*^*-/-*^ 1-3 days-old pups dissected in Ca^2+^/Mg^2+^-free Hank’s Balanced Salt Solution (HBSS, Invitrogen) and processed following standard procedures [79]. Cells were plated in Dulbecco’s modified Eagle medium and F-12 nutrient mixture (DMEM/F12, Invitrogen) containing 10% FCS (Invitrogen), and gentamycin (Sigma Aldrich). Cortices from two pups were platted in a 75cm^2^ flask pre-treated with Poly-D-Lys (P7280, Sigma Aldrich) and cultured at 37°C in a humidified 5% CO_2_ incubator. After 24h, the medium was refreshed and supplemented with m-CSF1 to improve microglial survival. Cultures were let reach confluency (about 2 weeks) without further changes. For mixed culture analysis, cells were detached by washing with warm PBS and then incubated with 0.25% trypsin (Invitrogen), 1mM EDTA in PBS at 37°C for 15min. After adding DMEM/F12 with 10% FCS, cells were centrifuged for 5 min at 1000 rpm, and re-suspended in the same medium. Microglial cells were purified by mechanical detachment of mixed glial cultures, recovered by medium centrifugation (5 min; 1000 rpm) [79] and plated replacing only 50% of the medium to promote microglial proliferation. Cells were seeded on multi-well culture plates (Falcon) at a density of 10^5^ cells/cm^2^. In mixed cultures, the astrocyte/microglia ratio was 1:1. Cells were let settle for 48h and treated with either LPS (1µg/ml) or saline. Cell media were then collected and processed.

### Immunohistochemistry

Newborn and adult mice were perfused transcardially with 4% PFA in PB 0.1M (wt/vol). Brains were removed, post-fixed by immersion overnight and then washed for 24h in PBS, on a rocking platform at 4°C. Spinal cords were extracted after body post-fixation and washing. Tissues were incubated in a 30% sucrose-PB solution (wt/vol) for 24h, embedded in a 7.5% gelatin; 15% sucrose solution (wt/vol), frozen on dry ice and, if necessary, stored at −80°C until serially sectioned coronally at 15µm of thickness using a cryostat (Leica). Histological and cytological immunostaining was performed following standard protocols, after antigen retrieval (10mM citrate buffer, pH6, for 5min at 110°C in a boiling chamber, Biocaremedical). Primary antibodies (Table S2) were incubated ON at RT. The following secondary antibodies were incubated for 1h at RT: Alexa488 or Alexa594-conjugated donkey anti-rabbit or anti-mouse (1:1000); goat anti-rat (1:3000; Molecular Probes, Invitrogen) or anti-chicken, (1:2000, AbCAM); biotin conjugated goat anti-mouse or anti-rabbit (1:500, Jackson Lab). Alexa488, Alexa594 (1:500; Molecular Probes, Invitrogen) or POD conjugated (Jackson Lab) streptavidin followed by reaction with 3,3-diaminobenzidine (0.05%; Sigma) and 0.03% H_2_O_2_. For immunofluorescence, sections were counterstained with Hoechst (Sigma Aldrich). Tissue was analysed with a DMCTR5000 microscope equipped with a DFC350Fx monochrome camera or a DFC500 color camera (Leica Microsystems) or with a LSM710 confocal imaging system coupled to an AxioImager.M2 vertical microscope (Zeiss) or a LSM800 coupled to an Axio Observer inverted microscope (Zeiss). Fluorescence was quantified with ImageJ software (National Institute of Health) using 12-14 sections per analyzed brain.

### ELISA

The amount of cytokines’ present in the glial culture media was determined with electro-chemo-luminescence in a MSD MULTI-SPOT Assay System using 96 well V-PLEX plates for pro-inflammatory mouse panel 1 or custom mouse cytokine V-PLEX plates for IFNγ, IL1β, IL4, IL6, IL10 and TNFα (Meso Scale Discovery), following the manufacturer’s indications and using a SECTOR Imager 2400 reader (Meso Scale Discovery). SFRP1 presents in glial culture medium or in the RIPA fraction of brain lysates was determined with a capture ELISA [23]. Culture media were diluted five folds, and brain lysates used at 0.1 µg/µl protein concentration determined with BCA protein assay (Thermo Scientific). Values were determined at 450 nm wavelength using a FLUOstar OPTIMA micro-titter plate reader (BMG LABTECH).

### Fluorescence Activated Cell Sorting (FACS) and Flow Cytometry Analysis

*CX*_*3*_*CR1*^*+/GFP*^ and *CX*_*3*_*CR1*^*+/GFP*^;*Sfrp1*^*-/-*^ 10-12 weeks-old male littermates were treated with LPS or saline as described. After 3 days, animals were perfused with ice-cold saline and brains collected on ice-cold, HBSS Ca^2+^, Mg^2+^-free (Invitrogen). After meninges’ removal, cortices were isolated, finely chopped and digested for 20 min in DMEM with GlutMAX (Invitrogen) containing papain 20U/ml (Worthington Biochemical Corporation), DNase (50U/ml, Sigma Aldrich) and L-Cysteine (1mM, MERCK). After addition of 20% FCS (Invitrogen), the tissue was mechanically dissociated and filtered through a 35µm nylon strainer (Falcon). Microglial cells were separated from myelin/debris by Isotonic Percoll gradient centrifugation (35% Fisher Scientific) at 2000 rpm for 45min at 4°C. The pellet was recovered and sequentially incubated with anti-mouse CD16/CD32 (1:250, BD Pharmingen; for 15min at 4°C) followed by rat anti-CD11b and PerCP-Cy5.5 rat anti-mouse CD45 (1:200, BD Pharmingen; for 30min at 4°C), all in 2% BSA, 5mM EDTA in PBS. After washing, cells were sorted using a BD FACS Aria Fusion Flow Cytometer and their signal, size and complexity acquired with DiVA8 Software (BD Pharmingen). Analysis was performed using FlowJo v10.0.7 Software (BD Pharmingen).

### Genomic Data Processing and Access

RNA from sorted microglia was extracted with the TRI reagent (MERCK) according to the manufacturer’s instructions, purified with RNeasy Lipid Mini Kit (Qiagen) following the manufacturer’s protocol and the resulting total RNA was treated with RNase-Free DNase Set (Qiagen). RNA quality was assessed with a Bioanalyzer 2100 system (Aligent), obtaining RIN (RNA integrity number) values between 8.8 and 10. Each sample (microglia sorted from a single mouse brain) was processed to obtain a RiboZero Stranded Gold Library (Illumina) and sequenced in HiSeq 4000 sequencer in paired-end configuration with 150bp sequence reads (Illumina). Sequence quality was determined with FastQC (Babraham Bioinformatics) revealing more than 32 million mean cluster reads of over 38 quality score and 93% mean Q30. Reads were aligned with HISAT2 v2.1.0 [80] to Ensemble *Mus musculus* (GRCm38.94). Aligned reads were further processed using Samtools v1.9 [81] and quantified to gene level using HTseq v0.11.2 [82]. Whole genome alignments were visualized with Integrative Genomics Viewer (IGV v2.5) [83]. Differential expression analysis (DGE) was performed using the Bioconductor package DESeq2 v1.10.0 [84]. DGE data were processed with custom R scripts (R version 3.5.1, 2018) considering genes with adjusted p-value < 0.05 and log2 Fold Change > +/-0.8 as significantly up-or down-regulated. Analysis of GO terms was performed with the Bioconductor package clusterProfiler [85] and that of promoter-based motif enrichment with HOMER [86]. RNAseq data sets can be accessed at the European Nucleotide Archive public repository under the following accession numbers (ERP119668 / PRJEB3647).

### Statistical analysis

was performed using Prism v7 software (GraphPad). Different statistical tests were used as indicated in each figure footnote and represented by *P<0.05, **P<0.01, ***P<0.001, ****P<0.0001.

## Acknowledgements

We wish to thank J Avila, CBM, for providing a breeding pair of the *CX*_*3*_*CR1::GFP* mice and the CBM image analysis, genomic and cytometry facilities for their support and advices during the course of this study. This work was supported by grants from the Spanish AEI (BFU2013-43213-P; BFU2016-75412-R with FEDER support), Fundación Tatiana Perez de Guzman el Bueno and CIBERER to PB and PE. JRC (BES-2011-047189) and MIM (BES-2014-068797) were supported by FPI fellowships from the MINECO. We also acknowledge a CBM Institutional grant from the Fundación Ramon Areces.

## Author contributions

JRC, PE and PB conceptualized and designed the research study. JRC, MIM and MJMB conducted the experiments and acquired the data. AB, JRC and BA designed, the EAE study and JRC and AB conducted and analyzed it. JRC, MIM, MTH, PE and PB analysed the data. MPK and SS provided support with cytokine analysis. JRC and PB wrote the paper. JPLA supervised RNAseq analysis. All authors read and approved the manuscript.

## Data Availability Section

Materials will be available upon request

RNAseq data can be accessed at the European Nucleotide Archive public repository: ERP119668 / PRJEB3647.

## Supplementary Figure Legends

**Supplementary Figure 1. SFRP1 enhances glial cell activation**. A) Confocal image analysis of cryostat sections from adult *Sfrp1*^*+/βgal*^ mouse brains three days after intra-cortical infusion of LPS. Sections were immunostained for βgal (green) and GFP (red) or Iba1 (red). Note that astrocytes but not microglial cells express the βgal/Sfrp1 reporter. B) On the left, diagram of the observed GFP distribution (green dots) after lentiviral (LV) particles’ delivery into the lateral ventricle (*Lv*). Images show examples of immunostaining against GFP protein (GFP signal was no longer visible, unless detected with immunohistochemistry) in the Lv of brains transduced with LV-*IRES-Gfp* or LV-*Sfrp1-IRES-Gfp* 5 months (m) post-infusion. C) Coronal sections from LV-*IRES-Gfp* or LV-*Sfrp1-IRES-Gfp* infected brains 1 or 5 months (m) post-infusion (PI). Sections were immunostained for CD45. Arrowheads indicate CD45^hi^-positive macrophages or lymphocytes infiltrated in the parenchyma after prolonged LV-*Sfrp1-IRES-Gfp* infection. High power images 1 and 2 were taken from the regions indicated with grey dotted lines in A. D) Coronal sections from wt and *Sfrp1*^*-/-*^ mouse brains three days after infusion of saline or LPS, immunostained for GFAP. The white arrows indicate the injection site; the area boxed with white lines is represented at high power in Fig 3.

**Supplementary Figure 2. Astrocytes-derived SFRP1 enhances glial cell activation upon LPS treatment**. A) The graph shows the % CD11b+ populations present in the cortex of *CX*_*3*_*CR1*^*GFP/+*^ or *CX*_*3*_*CR1*^*GFP/+*^;*Sfrp1*^*-/-*^ mice 3 days after saline or LPS intraventricular infusion. Gating was set for isolating the following myeloid cell populations: CD11b^+^;CD45^lo^;GFP^+^ surveying microglia, CD11b^+^;CD45^+^;GFP^+^ activated microglia and CD11b^+^;CD45^+^;GFP^-^ infiltrated monocytes. B) ELISA determination of SFRP1 levels present in the media of microglia or microglial and astrocytes culture derived from wt or *Sfrp1*^*-/*-^ pups. Note that SFRP1 is detected only in cultures in which astrocytes are present. Note also that LPS induces a significant increase of secreted SFRP1 protein. Error bars represent Standard Error. Statistical significance: ****P<0.0001; One way ANOVA followed by Bonferroni Test

**Supplementary Figure 3. Gating strategy to assess phagocytosis**. A) Representative cytometry plots of brain cells suspension showing gating of individualized live cells CD11b^+^;CX3CR1^+^ recognized as microglial cells for pHrodo labeled *E. coli* bio-particles quantitation. Comparison of microglial cells exposed (*E*.*coli* Ctrl+) or not exposed (*E*.*coli* Ctrl-) to pHrodo labeled *E*.*coli* shows specific pHrodo recognition. B) Representative confocal image of FACS sorted GFP^+^ microglia. Note specific pHodo^+^ lysosomal inclusion of *E*.*coli* particles in phagocyting vs non-phagocyting microglia.

**Supplementary Figure 4. SFRP1 microglial inflammatory response depends on HIF factors**. A) Representation of the four up-regulated k-means unsupervised clusters of up-regulated genes in response to LPS in *CX*_*3*_*CR1*^*GFP/+*^ or *CX*_*3*_*CR1*^*GFP/+*^;*Sfrp1*^*-/-*^ microglial cells. Eight different clusters were generated by Z-scores co-variation. The clusters are indicated in the heatmap of Fig. 5F. B) Gene Ontology Enrichment Analysis of different clusters, GO biological process annotated are represented by fold enrichment and color-coded by adjusted p-value of enrichment and number of genes of each cluster in that term. C) Regulatory Elements Analysis of different clusters. Specifically enriched transcription factors are represented by fold enrichment relative to the overall transcripts present in the microglia transcriptome. Colored by Log odds detection threshold and the percentage of total targets for each transcription factor present in the cluster.

