## Supplementary figures and images for "SFRP1 shapes astrocyte to microglia cross-talk in acute and chronic neuroinflammation"

### Fig S1

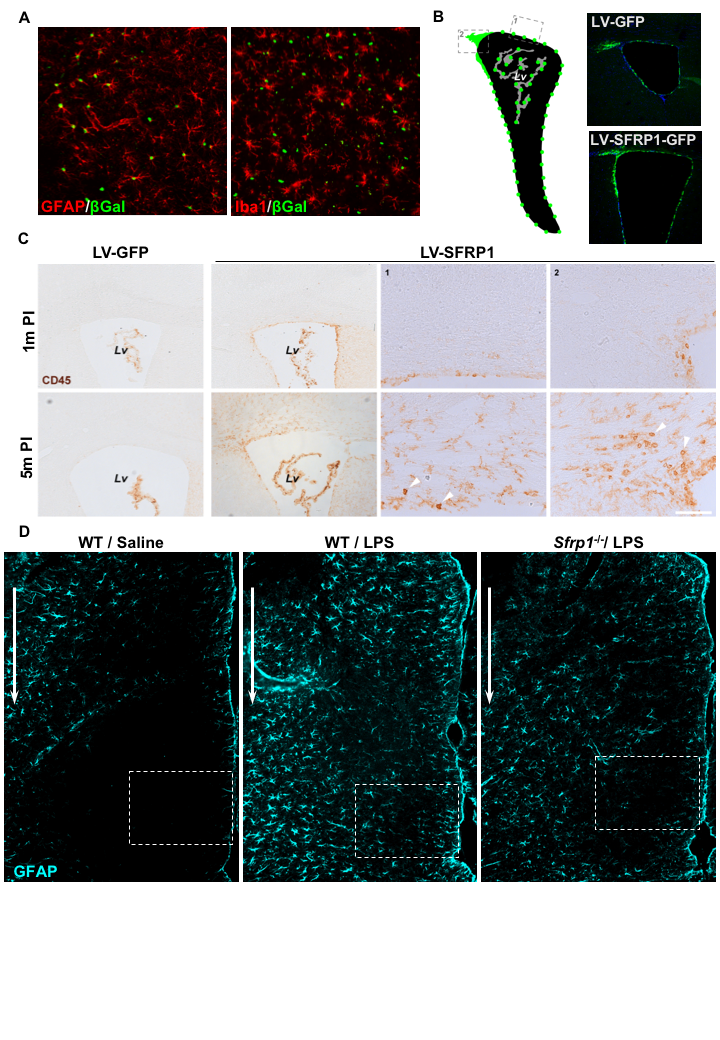

### Fig S2

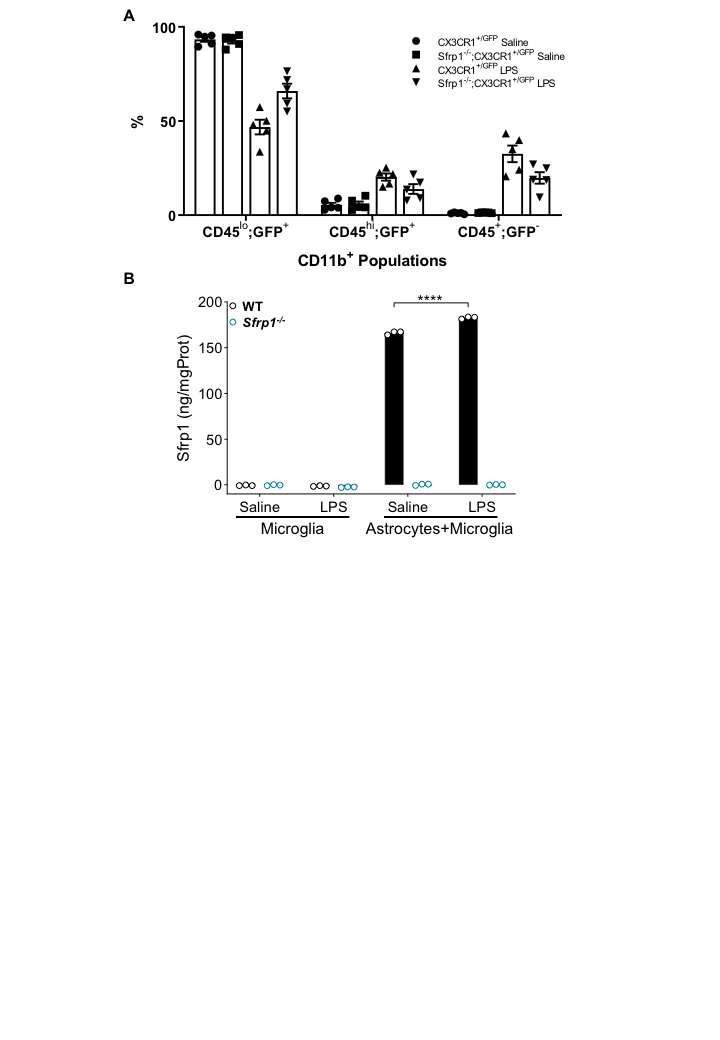

### Fig S3

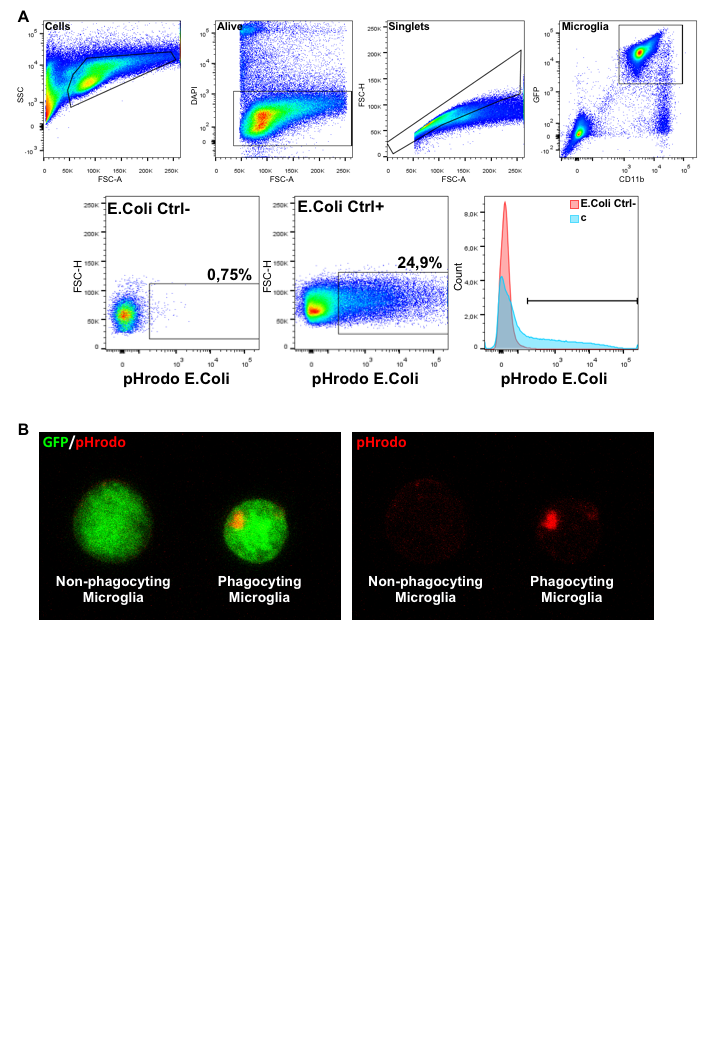

### Fig S4

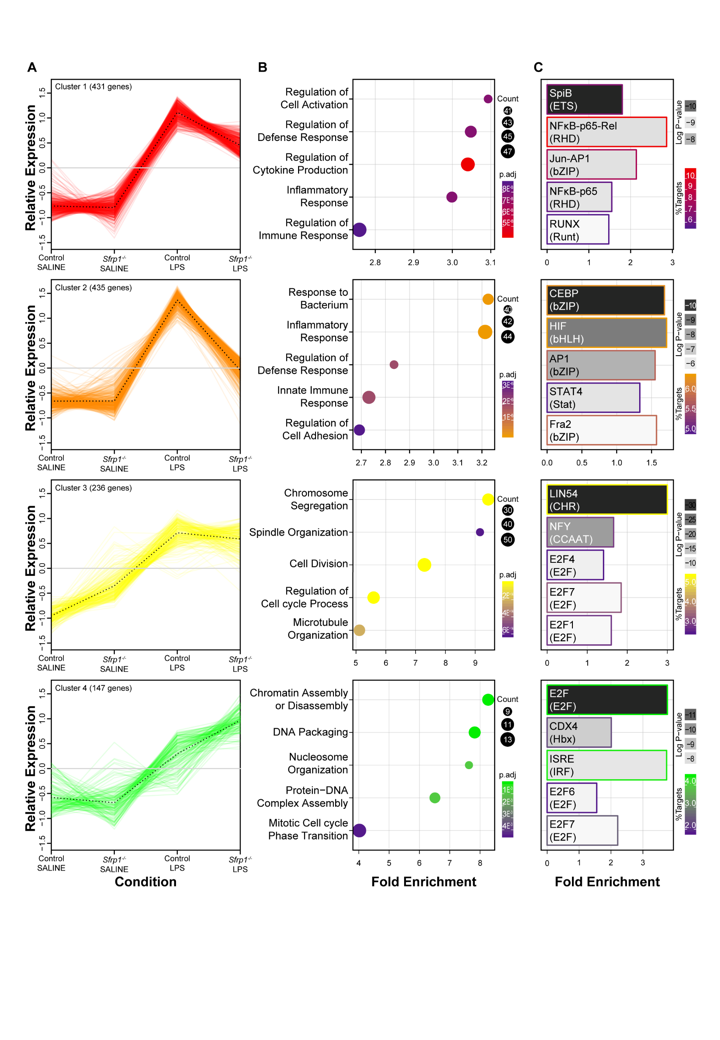
